# Quantification of water exchange across the blood-brain barrier using non-contrast MR fingerprinting

**DOI:** 10.1101/2023.11.15.567199

**Authors:** Emma L. Thomson, Elizabeth Powell, Claudia A. M. Gandini Wheeler-Kingshott, Geoff J. M. Parker

**Affiliations:** Centre for Medical Image Computing, Department of Medical Physics and Biomedical Engineering, University College London, London, United Kingdom; NMR Research Unit, Queen Square MS Centre, Department of Neuroinflammation, UCL Queen Square Institute of Neurology, Faculty of Brain Sciences, London, United Kingdom; Department of Brain Behavioural Sciences, University of Pavia, Pavia, Italy; IRCCS Mondino Foundation, Pavia, Italy; Bioxydyn Limited, Manchester, United Kingdom

**Keywords:** Blood brain barrier, MR fingerprinting, water exchange

## Abstract

**Purpose:** A method is proposed to quantify cerebral blood volume (*v*_*b*_) and intravascular water residence time (*τ*_*b*_) using magnetic resonance fingerprinting (MRF), applied using a spoiled gradient echo sequence, without the need for contrast agent.

**Methods:** An in silico study optimised an acquisition protocol to maximise the sensitivity of the measurement to *v*_*b*_ and *τ*_*b*_ changes. Its accuracy in the presence of variations in *T*_1,*t*_, *T*_1,*b*_, and *B*_1_ was evaluated. The optimised protocol (scan time of 19 minutes) was then tested in a exploratory healthy volunteer study (10 volunteers, mean age 24 ± 3, 6 male) at 3 T with a repeat scan taken after repositioning to allow estimation of repeatability.

**Results:** Simulations show that assuming literature values for *T*_1,*b*_ and *T*_1,*t*_, no variation in *B*_1_, while fitting only *v*_*b*_ and *τ*_*b*_, leads to large errors in quantification of *v*_*b*_ and *τ*_*b*_, regardless of noise levels. However, simulations also show that matching 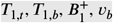 and *τ*_*b*_, simultaneously is feasible at clinically achievable noise levels. Across the healthy volunteers, all parameter quantifications fell within the expected literature range. In addition, the maps show good agreement between hemispheres suggesting physiologically relevant information is being extracted. Expected differences between white and grey matter *T*_1,*t*_ (p<0.0001) and *v*_*b*_ (p<0.0001) are observed, *T*_1,*b*_ and *τ*_*b*_ show no significant differences, p=0.4 and p=0.6 repectively. Good repeatability was seen between repeat scans: mean ICC of *T*_1,*t*_ : 0.91, *T*_1,*b*_ : 0.58, *v*_*b*_ : 0.90, and *vτ*_*b*_ : 0.96.

**Conclusion:** We demonstrate that regional simultaneous quantification of *v*_*b*_, *τ*_*b*_, *T*_1,*b*_,*T*_1,*t*_, and *B*_1_ using MRF is feasible in vivo.

## 1 INTRODUCTION

Integrity of the blood brain barrier (BBB), the semi-permeable barrier that separates the blood in vessels from the extracellular tissue in the central nervous system (CNS)^1^, is vital for supplying the brain with nutrients and solutes while protecting the neural tissue from toxins and pathogens. Break-down of the BBB allows these toxic substances into the brain and is thought to be a key component in the progression of multiple neurological diseases.^2^

The ability to quantify physical parameters associated with BBB integrity would give insight into early disease and may be valuable for monitoring progression and response to treatment. A number of solutions for the detection of break-down of the BBB using MRI have been proposed; currently the most widely available method, DCE-MRI^3,4^, requires a gadolinium based contrast agent (GBCA), and is best suited to quantifying severe damage, as it measures the transfer of the relatively large contrast agent complex over the BBB. Aside the fact that GBCA carries some health-related contraindications^5^, it suffers from poor sensitivity and a low signal-to-noise ratio (SNR) when trying to assess subtle damage.^6,7^

An alternative approach, which may offer greater sensitivity than DCE-MRI, while also avoiding the use of GBCAs, is to measure exchange of the smaller, and endogenously abundant, water molecules across the BBB. However, existing water exchange techniques, such as those based on arterial spin labeling (ASL)^8,9,10^ or diffusion MRI methods such as filter echange imaging (FEXI)^11,12^, also suffer from low SNR.

A technique that has been shown to boost SNR and sensitivity in a range of quantitative MRI settings is magnetic resonance fingerprinting (MRF). MRF exploits the signal response of tissues when repeatedly exposed to radiofrequency (RF) pulses of different amplitudes at varying intervals.^13,14,15^ Multiple sequence repetitions are performed consecutively, generating a signal that is characteristic of a unique set of tissue properties. These voxel-wise responses are compared with entries in a dictionary of simulated responses calculated from a set of known parameters, allowing maps of multiple parameters to be extracted simultaneously and efficiently.

Here, we propose, for the first time, using MRF to quantify BBB water exchange. We propose a technique that uses an RF spoiled gradient echo (SGRE)-MRF sequence to enable the simultaneous quantification of cerebral blood volume (*v*_*b*_), intravascular residence time (*τ*_*b*_, the inverse of water exchange rate), intravascular *T*_1_ (*T*_1,*b*_), extravascular *T*_1_ (*T*_1,*t*_), and *B*_1_ multiplication factor, 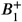, without the use of contrast agent.

Through an in silico study an acquisition protocol was optimised. The accuracy and sensitivity of the measurement in the presence of variations in intravascular *T*_1_, extravascular *T*_1_, and 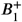 was evaluated. The optimised protocol was then tested in a healthy volunteer study to determine feasibility and repeatability.

## 2. METHODS

### 2.1 Simulations and Optimisation

A 3D array of spin isochromats was simulated assuming a two-site exchange system between intra and extravascular pools, with defined compartmental volumes and a semipermeable barrier, representing the BBB, allowing for two-way exchange. A singular, spatially non-uniform, voxel was modelled, described by a grid of isochromats in the x-y-z plane, visualised in Figure 1 . The Bloch equations were used to simulate signals arising from this array and its magnetisation evolution over time. The isochromats within the intravascular compartment had a residence time, *τ*_*b*_, dictating how long on average they spent in the vasculature. Exchange was simulated by calculating the cumulative probability of exchange for each isochromat at each simulation time step. Diffusion was assumed to occur freely within each compartment so the exchanging isochromat from the intravascular compartment would exchange with a random isochromat from within the extravascular compartment. Volume fractions were considered constant, so all exchange is two way: exchanging isochromats “swap” compartments while retaining magnetization history. After exchange, the cumulative probability of exchange for each exchanging isochromat is reset to zero. Simulated signals were generated first in quadrature and then summed such that the resultant transverse magnitude data was stored as the measured signal.

**FIGURE 1.**
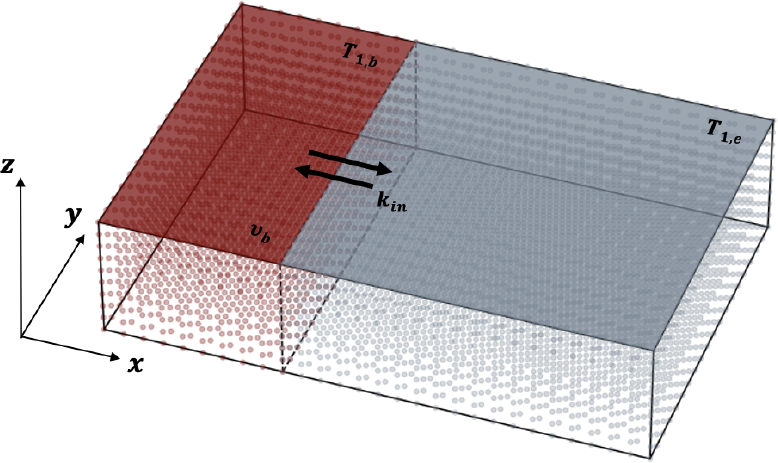
Grid array of isochromats subject to Bloch equation magnetization changes, in which a percentage (*v*_*b*_) are governed by *T*_1,*b*_ (the blood compartment in red) and the remainder are governed by *T*_1,*t*_ (the tissue compartment in grey). Limited exchange occurs between the two compartments with the probability of exchange governed by the mean intravascular residence time (*τ* _*b*_ = 1/*k*_*in*_) . Exchange influx and efflux is assumed to be equal. Along the *z* dimension a variation in flip angle magnitude was applied to reflect the slice profile.

Along the z-direction the applied flip angle varies according to the imaging sequence slice profile. Slice profile was calculated using a separate Bloch equation simulation^16^ using the known characteristics of the applied RF pulse. To reduce the number of isochromats simulated, the slice profile was assumed to be symmetrical so only half of the profile was sampled and the resultant signal magnitude multiplied by two.

Optimisation was performed using a branch and bound algorithm^17^ such that the variations of *α* and *TR* were optimised to maximise the difference between signals at the extremes of the range of healthy *τ*_*b*_ values (200 ms ≤ *τ*_*b*_≤1600 ms).^8^ The optimal set of sequence parameters were the ones that maximised the inner product between the two signals. *T*_1,*t*_ was fixed at 1300 ms^18,19^, and *T*_1,*b*_ at 1700 ms^20^, mimicking a grey matter voxel at 3 *T*. Variation of *α* and *TR* was limited to predetermined shapes, with amplitude and width allowed to be optimised to allow for a computationally feasible search space.

The flip angle was varied to give a periodic positive sinusoidal variation in which the even peaks were lower than the odd, emulating the flip angle design in the original MRF paper^13^, using the equation:

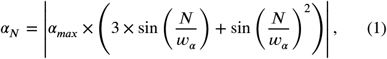

where *N* is the *N*^th^ repetition of the applied sequence, and *α*_*max*_ and *w*_*α*_ are the parameters that govern the maximum height of the peaks (degrees) and the width of the periodic variation (# of *TR*s), respectively. Optimisation bounds on these parameters were (20 ≤*α*_*max*_≤88) [degrees] and (5 *π* ≤ *w*_*α*_ ≤ 200 *π*) [# of *α*].

The variation in *TR* was sinusoidal governed by:

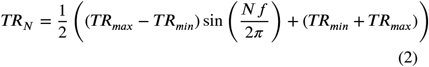

where *f* is the frequency that governs the period of the sinusoidal variation [(# of *α*)^−1^], and *TR*_*min*_ (ms) and *TR*_*max*_ (ms) govern its range. Optimisation limits for these parameters are: (1 ≤*T*≤200) [(# of *α*)^−1^], (4 ≤ *TR*_*min*_ ≤200) [ms] and (9 ≤*TR*_*max*_ ≤ 200) [ms]. An upper boundary was imposed on the possible repetition time such that a 2000 *TR* sequence with a fully-sampled centre of k-space could be acquired in a clinically feasible times (For a scan time of less than 20 minutes, *TR*_*max*_ = 200 ms). *TR* lower bound was dictated by scanner sequence implementation lower limits (*TR*_*min*_ = 4 ms). These variation parameters can be visualised in Figure 2

**FIGURE 2.**
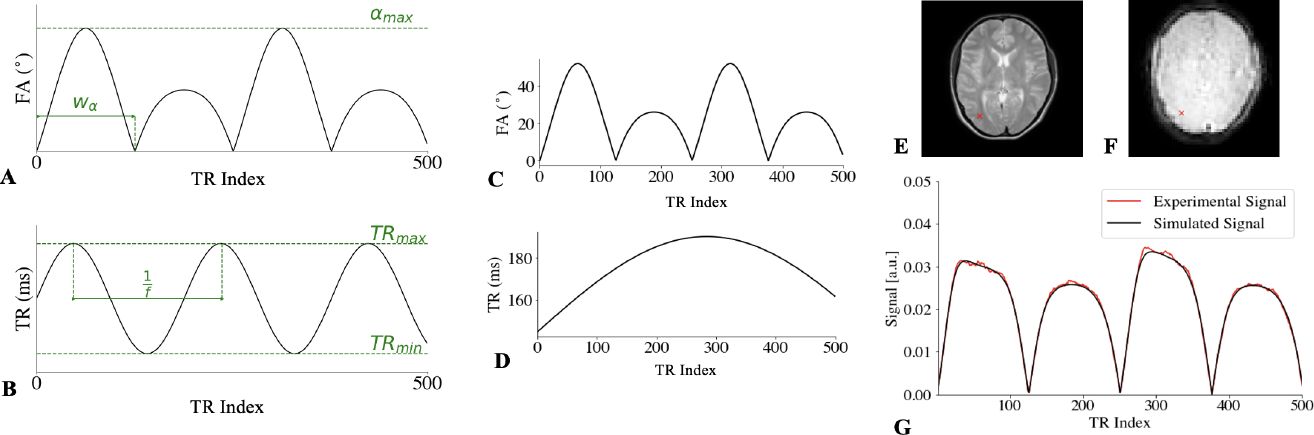
(a) Representative plots of the variations of the flip angle and (b) repetition time over five hundred consecutive repetitions of the sequence. The optimisable parameters that modify these variations are indicated in green. (c) Partial visualisation of optimal variation in flip angle (*α*) and (d) repetition time (*TR*) used for the first 500 acquisition steps: this variation repeats for the remaining *TR*s. The signals generated by these variations are matched to a precomputed dictionary. An example voxel marked in red on (e) a *T*_2_-weighted anatomy scan and (f) a raw MRF image (image 100/2000) was isolated. (g) Shows this voxel signal plotted against its closest dictionary match. 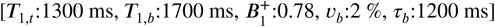

Optimisation was performed three times with 2000 *TR*s: the best of these three repeats was chosen to be the optimum set of parameters.

To test these simulations three sets of experiments were performed. For each experiment a dictionary was generated with a range of noise levels to test the matching sensitivity to noise: this was repeated 50 times for each dictionary to allow for statistical analysis of the quality of the match to be performed. Quantification was said to failed when the mean difference in matched value from the ground truth was larger than half a dictionary step size.

A 2-dimensional (2D) dictionary was generated with variation in blood volume from 1 to 10% in steps of 1%, denoted *v*_*b*_ = [*min* = 1 : *step* = 1 : *max* = 10]% and mean residence time of water in the intravascular compartment, *τ*_*b*_ = [200 :100 : 1600] ms. Three experiments were then performed: first, this 2D dictionary was matched against a 2D data set with the same variation to test the sensitivity of these variations to noise. Next, a sample data set with variation along 5 dimensions (5D), *v*_*b*_ ([1 : 1 : 10]%), *τ*_*b*_ ([200 : 100 : 1600] ms), intravascular *T*_1_: *T*_1,*b*_ ([1500 : 200 : 1900] ms), extravascular *T*_1_: *T*_1,*t*_ ([1000 : 200 : 2000] ms), and relative 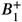: ([0.8 : 0.1 : 1.2]), was matched to the 2D dictionary containing variation in *v* _*b*_ and *τ*_*b*_ only, with fixed values of *T*_1,*b*_, *T*_1,*t*_, and 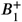 from the literature, to explore robustness of matching when these parameters are not known *a priori*. Finally, the 5D sample was matched to a 5D dictionary with the same variation as the sample data set to test the feasibility of determining each parameter simultaneously. Variations in each parameter were chosen to encapsulate healthy ranges for both white and grey matter.^8,21,18,19,22,23,24,25,20,26,27,28^.

#### 2.1.1 Noise Metric

We modelled noise as a zero-mean complex Gaussian with standard deviation (*σ*_*G*_ on each isochromat. Each isochromat has an equilibrium magnetization of unity such that (*σ*_*G*_ = 0.01 is equivalent to 1% noise. MRF signals however, seldom recover close to the equilibrium value: across a coarse, broad scope dictionary, the average signal value was 13.04% of the equilibrium value. For this MRF implementation we therefore define the signal-to-noise ratio as:

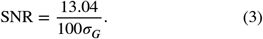

### 2.2 Healthy Volunteer Testing

#### 2.2.1 Acquisiton

Ten healthy volunteers (19-29 years old, mean age: 24 ± 3 years, 6 male) were recruited in accordance with the local Institutional Review Board guidelines with informed written consent. Each individual was scanned at 3 T on a Philips Ingenia system using a T1-fast field echo (T1-FFE, RF-spoiled gradient echo) acquisition with an MRF sequence of 2000 *TR*s, FOV = 224 mm x 244 mm, reconstructed matrix = 64 × 64, slice thickness = 5 mm, *TE* = 2 ms, fully sampled variable density spiral readout, SPIR fat saturation, with variation in *α* and *TR* as outlined in Figure 2, scan time = 19:27. A single central axial slice was acquired. A higher resolution single slice *T*_2_-weighted (Turbo Spin Echo, FOV = 224 mm x 244 mm, reconstructed matrix = 560 × 560, slice thickness = 5 mm, TE = 80 ms, scan time = 1:48) and 3D *T*_1_-weighed image (T1-FFE, FOV = 240 mm x 240 mm, reconstructed matrix = 256 × 256, slice thickness = 5 mm, *TE* = 1.854 ms, number of slices = 55, scan time = 3:22) were also acquired at the same location for segmentation and analysis. A repeat scan was performed within the same session as the initial scans after repositioning of the subject to allow assessment of parameter scan-rescan repeatability.

#### 2.2.2 Preprocessing

Brain extraction was done using FSL BET^29^, an initial affine linear registration performed using FLIRT^30,31^, a subsequent non-linear registration using FNIRT^32,33^, and segmentation performed using SynthSeg.^34^ As large voxel sizes lead to partial volume effects, partial volume segmentations for white and grey matter were thresholded so that only voxels that contained 90% of the respective tissue were used in the calculation of regional means.

#### 2.2.3 Dictionary and Matching

A 5D dictionary with around 0.65 million entries was generated for matching to experimental data. It consisted of variation along *T*_1,*b*_ ([1500 : 100 : 1900]ms), *T*_1,*t*_ ([600 : 50 : 2000]ms), *v*_*b*_ ([1 : 1 : 10]%), *τ*_*b*_ ([200 : 100 : 1600]ms) and 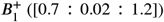. In addition to this an additional *T*_1,*t*_ value of 3000 ms was included to account for CSF values that may still be present post masking for partial volume effects. Matching was performed using the inner-product method^13^.

Matching of 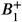 and *T*_1_ simultaneously poses a challenge in MRF due to difficulties in separating differences in signal due to *T*_1_ changes from those due to 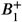 changes when the signal is normalised.^35,36,37^ Erroneously high matched 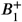 values are reported around the centre of the brain, visible in Figure 3 : this effect is prominent at higher *B*_1_ values. These ‘spikes’ in 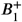 have a knock on effect on the remaining parameters. This was alleviated by performing a multi-stage matching process and utilising the knowledge that 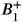 varies smoothly across the image. After an initial matching iteration using the 5D dictionary was performed, the matched 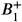 map underwent the following steps: First, it was measured in initially matched 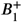 maps that the corrupt values all fell above 1.06. Therefore, all matched values that fell above 1.06 were considered to be corrupted. The corrupted high 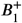 values were then removed along with voxels corresponding to CSF and the remaining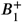 map fitted by triangulating the input data^38^ and constructing a piecewise cubic interpolating Bezier polynomial on each triangle^39^. This interpolation was repeated iteratively until the 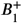 map was fully restored, Figure 3 . Upper and lower bounds were imposed on the interpolation to ensure that the process was performing correctly: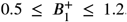 . The resultant map was smoothed with a Gaussian filter (*σ* = 1). This new 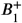map was re-discretised to the coarseness of the dictionary and fixed for each voxel. A second match was then performed with fixed 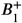 value.

**FIGURE 3.**
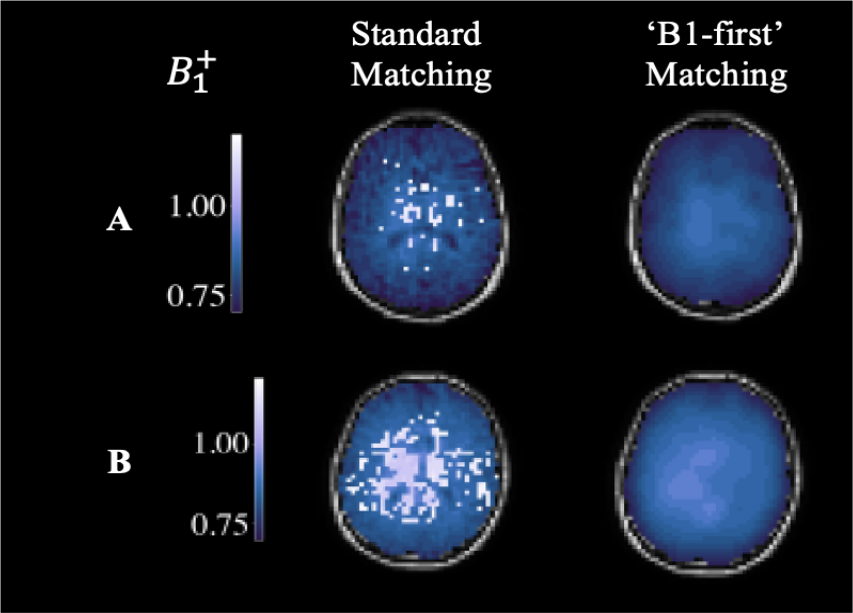
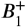 maps quantified simultaneously with *T*_1_ using the standard matching technique^13^ and the proposed 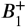 matching technique. For a) a volunteer with a low number of erroneous values and b) a volunteers with a large corrupted area. ‘*B*_1_-first’ matching corrects these erroneous values. The quantification of the remaining parameters is then re-performed with a fixed 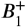 value to correct the corresponding voxels.

After matching, the median value across each segment identified by the SynthSeg segmentation was taken. Repeatability assessment involved generation of Bland-Altman plots of each parameter and calculation of their repeatability coefficient (RC).^40^

#### 2.2.4 Noise Metric

SNR for experimental results was defined as the mean value of the signal across the time course in each voxel divided by the residual sum of squares between the experimental signal and its closest dictionary match:

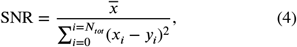

where *x* and *y* are the experimental signal and its closest dictionary match respectively, and *N*_*tot*_ is the total number of applied *TR*s. This provides a measure by which to compare simulated and experimental results.

#### 2.2.5 Statistical Analysis

Statistical analysis of the repeatability was performed using the repeatability coefficient (RC) and intra-class correlation coefficient (ICC). RC provides the expected difference in value between two repeats with a 95 % accuracy^41^, while ICC describes the resemblance between two groups scaled between 0-1.^42^ The scale described by Koo and Li (2016)^43^ was used to characterise ICC scores: less than 0.50 - poor, between 0.50 and 0.75 - moderate, between 0.75 and 0.90 - good, greater than 0.90 - excellent.

## 3 RESULTS

### 3.1 Simulation Experiments

The optimised values governing *α* and *TR* were found to be *α*_*max*_ = 52 degrees, *w*_*α*_ = 40*π* # of *TR*s, *f* = 181 (# of *TR*)^−1^, *TR*_*min*_ = 100 ms, and *TR*_*max*_ = 190 ms. A visualisation of these variations in *α* and *TR* can be seen in Figure 2 .

When implementing a 2D dictionary with no variation in *T*_1,*t*_, *T*_1,*b*_, or 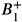 values, quantification was most robust to noise in the presence of large blood volumes and long residence times (low exchange), seen in Figure 5 A . When variation is present in *T*_1,*t*_ and *T*_1,*b*_, and *B*_1_, assuming literature values leads to large errors regardless of noise levels, Figure 5 B : neither *v*_*b*_ or *τ*_*b*_ are quantifiable under these conditions in a noise-free simulation, with a mean deviation from the ground truth in matching of 4.4 % and 715 ms for *v*_*b*_ and *τ*_*b*_, respectively.

**FIGURE 4.**
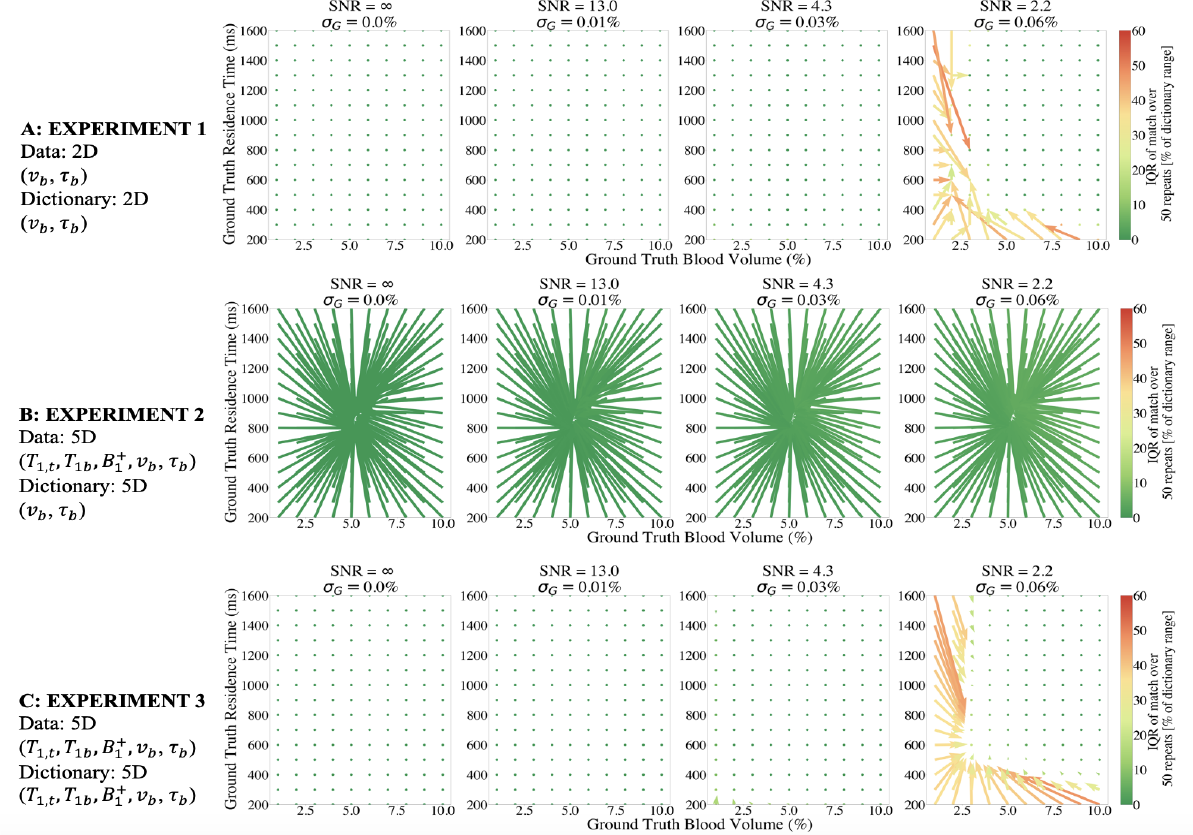
The success of matching a fingerprint to its corresponding dictionary entry for (A) *v*_*b*_ and *τ*_*b*_ with a fixed *T*_1,*b*_,*T*_1,*t*_, 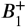, (B) *v*_*b*_ and *τ*_*b*_when there is discrepancy in the assumed values for *T*_1,*b*_,*T*_1,*t*_, 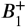, and (C) *v*_*b*_, *τ*_*b*_*T*_1,*b*_,*T*_1,*t*_ and 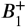 simultaneously. Each plot displays the 2D dictionary laid out with each residence time (*τ*_*b*_) and blood volume (*v*_*b*_) combination. Arrows indicate the deviation in matching from the ground truth value - the bottom of arrow is situated at the true value, the tip at median matched value. The interquartile range of match across 50 repeats is indicated by the colour of the arrow. No arrows indicates perfect matching. This can be seen across 4 example noise levels.

**FIGURE 5.**
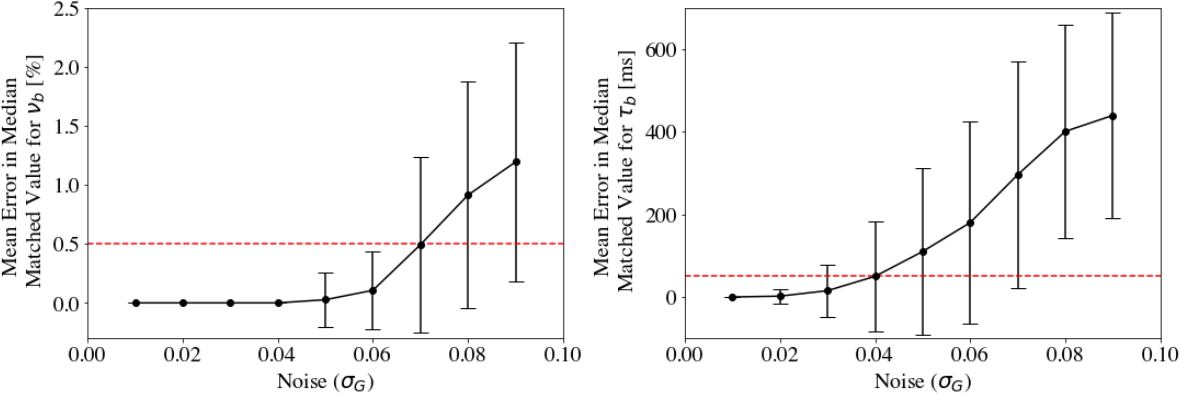
the success of quantifying a) *v* _*b*_ and b) *τ*_*b*_ across different noise levels. The matching is said to have failed when the mean difference in the median matched value from the ground truth is larger than half a dictionary step size: this threshold is marked by the dotted red line.

Using the 5D dictionary (Figure 5 C), results in less noise in the matched values than when using the 2D dictionary (Figure 4) across both *v*_*b*_ and *τ*_*b*_; this is due to averaging effects in the larger dictionary (the 5D dictionary is 90 times larger than its 2D counterpart). Comparing the matching success of *v*_*b*_ and *τ*_*b*_ between the 2D and 5D dictionaries we can see that there is no significant difference in accuracy (*σ*_*G*_ = [0.06, 0.09], *v*_*b*_ : *p* = [0.2745, 0.0562], *τ*_*b*_ : *p* = [0.1590, 0.2311]) or precision (*σ*_*G*_ = [0.06, 0.09], *v*_*b*_ : *p* = [0.1396, 0.2245], *τ*_*b*_ : *p* = [0.2360, 0.6370]) between the dictionaries . For this level of dictionary coarseness, no error is seen in the matching of *T*_1,*t*_, *T*_1,*b*_, or 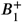 at these noise levels.

Figure 5 illustrates the SNR required to successfully quantify *v*_*b*_ and *τ*_*b*_ when using the 5D dictionary. Looking at the mean deviation of the matched value from the ground truth for both *v*_*b*_ and *τ*_*b*_ across the dictionary, and imposing a threshold of success of a mean deviation of half a dictionary step size or less, we can see that *v*_*b*_ can be quantified accurately up to a noise level equal to *σ*_*G*_ = 0.07 (corresponding to an SNR of 1.9), while the *τ*_*b*_ is only quantifiable up to a noise level of *σ*_*G*_ = 0.04 (SNR of 3.3).

### 3.2 Healthy Volunteer Testing

The above simulation results informed the quantification of the healthy volunteer data: voxels with an SNR of less than 3.3 were removed from calculations (mean SNR across volunteers was 11.86). The maps of removed voxels can be seen in Supplementary Information Figure 1.

Maps of *T*_1,*t*_, *T*_1,*b*_, 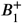, *v*_*b*_ and *τ*_*b*_ for a single volunteer alongside their repeat scan can be seen in Figure 6 . Maps for all volunteers can be seen in Supplementary Information Figure 2 . Maps of each parameter are visually consistent between repeats and between individuals. Regional variation is apparent in the maps of *T*_1,*t*_ and *v*_*b*_, while all other maps show little spatial variation. The patterns of spatial variation are largely consistent between scans for *T*_1,*t*_, while the spatial distribution of *v*_*b*_ is visually less repeatable.

**FIGURE 6.**
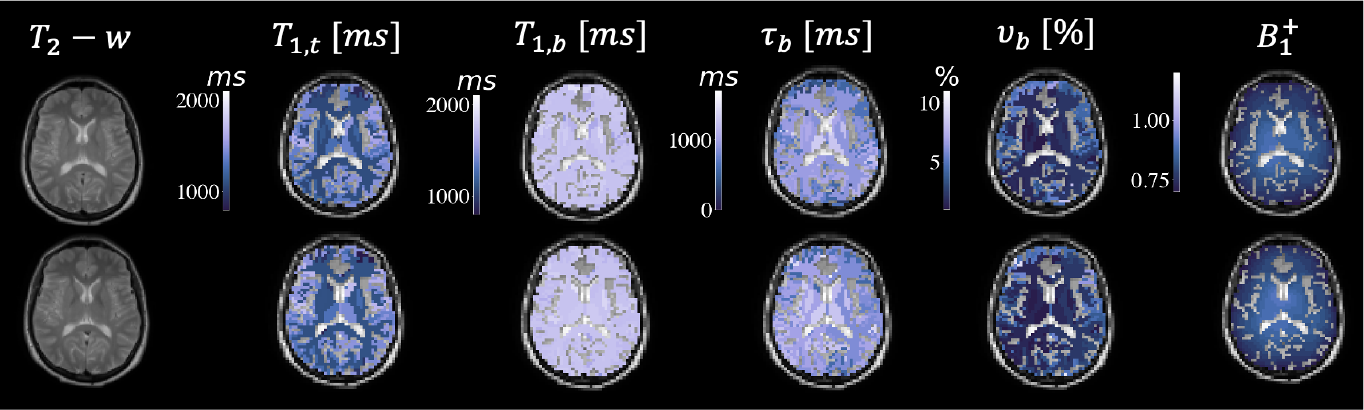
Regional quantitative parameter maps of *T*_1,*t*_, *T*_1,*b*_, *v* _*b*_ and *τ*_*b*_ for a single volunteer with repeat scan, and a voxel wise map for 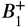.

The mean values for each parameter across the ten subjects can be seen in Figure 7 . Mean *T*_1,*t*_ differed between white and grey matter (1060 ± 50 ms and 1290 ± 70 ms, respectively (*p* < 0.0001)), as did mean *v*_*b*_ (1.9 ± 4 % and 2.8 ± 7 %, respectively (*p* < 0.0001)). *T*_1,*b*_ values were highly similar for white and grey matter (1750 ± 10 ms and 1756 ± 10 ms, respectively (*p* = 0.4270)), as were *τ*_*b*_ (930 ± 100 ms and 950 ± 150 ms, respectively (*p* = 0.6609)).

**FIGURE 7.**
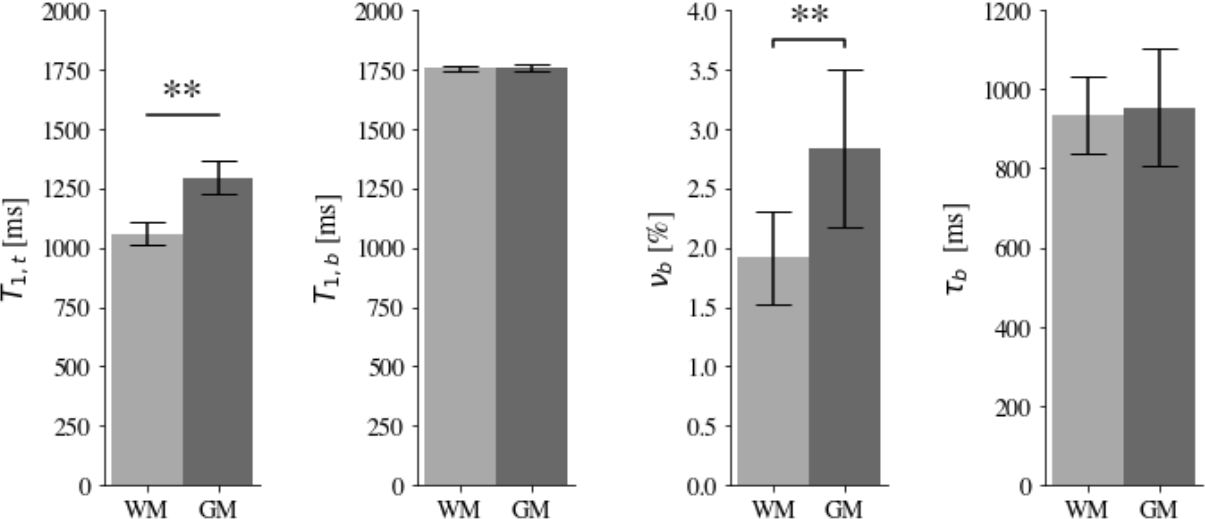
Global white and grey matter values averaged across all ten volunteers.

Bland-Altman plots (Figure 8) show the scan-rescan variation in mean parameter values for white matter and grey matter. Negligible bias is observed between measurements. Repeatability coefficients and intraclass correlation coefficients for each parameter are shown in Table (1). The ICC indicates high levels of repeatability for all parameters except *T*_1,*b*_ where the repeatability is moderate for grey matter and poor for white matter.

**TABLE 1.** The mean values, repeatability coefficient and intraclass correlation coefficient for white and grey matter across the four quantified tissue parameters.

|  | WM |  |  | GM |  |  |
| --- | --- | --- | --- | --- | --- | --- |
|  | Mean | RC | ICC | Mean | RC | ICC |
| $T_{1,t}$ [ms] | 1060 | 38.70 | 0.92 | 1290 | 67.59 | 0.89 |
| $T_{1,b}$ [ms] | 1750 | 22.01 | 0.56 | 1750 | 23.61 | 0.59 |
| $v_b$ [%] | 1.9 | 0.44 | 0.85 | 2.8 | 0.48 | 0.94 |
| $\tau_b$ [ms] | 930 | 45.74 | 0.97 | 950 | 108.75 | 0.94 |

**FIGURE 8.**
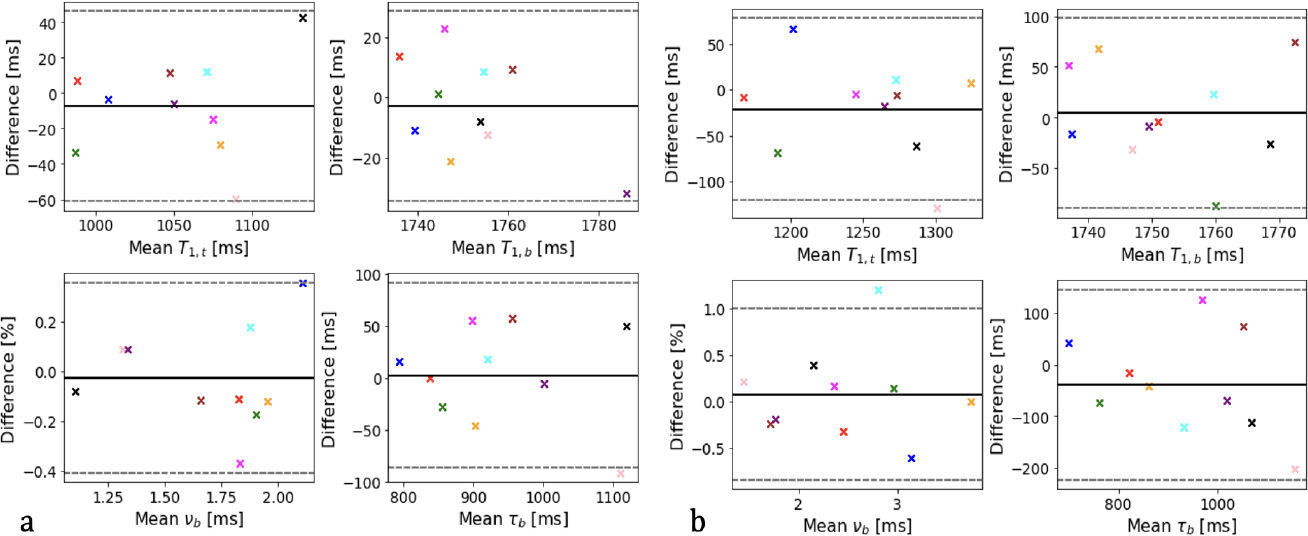
Bland-Altman plot comparing repeat measurements for *T*_1,*t*_, *T*_1,*b*_, 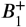 *v*_*b*_, and *τ*_*b*_ for (a) white matter, and (b) cortical grey matter. Marker colour is consistent across plots for each volunteer. Solid black line indicates mean difference between scan/rescan, with the 95% confidence intervals shown with dotted lines

## 4 DISCUSSION

### 4.1 Simulation Experiments

The simulation-based optimisation of the data acquisition focused on optimising the patterns of flip angle and repetition time variation. A parametric form was used to enable optimisation that included five parameters controlling the height and width of regularly shaped curves (Equations 1, 2 and Figure 2). This approach has the advantage of simplicity and ease of optimisation, but it limits the potential patterns of variation to those that can be parameterised using these simple functions. Other work has focused on a free-form optimisation of input parameters using the Cramer-Rao bound objective function.^44,45^ However, due to the high dimensionality of our optimisation problem, the computational and time requirements render this unfeasible.

Lack of success of quantification is seen when there is variation in *T*_1,*t*_, *T*_1,*b*_ and 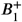 that is not accounted for. Inaccurately matched values of dictionary-based matching techniques inherently tend towards the average signal, in the case of a uniformly sampled dictionary, the central value. This suggests that, unless independent measures of these parameters are provided, all five parameters need to be quantified simultaneously using a 5D dictionary.

In order to reduce the dimensionality of the problem posed here an SGRE sequence was selected with a short *TE* = 2.1*ms*. This removes dependence of the signal on 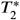, reducing the number of parameters that need to be quantified simultaneously.

The largest limitation of the underlying model (as with any model) is that the pathology of the human brain is far too complex to simulate in its entirety. This means that models can only account for a select few effects on signals, with the intention that these effects describe the meaningful majority of the observed signal variance; a simulation will never be completely accurate. Here, a two compartment generative model is assumed, which neglects, for example, exchange between plasma and red blood cells, and exchange between interstitial fluid and the various cells found in the brain. In addition it is assumed that water is well mixed in each compartment, with exchange being equally probable for spins throughout each compartment, rather than being more likely for those closer to the blood brain barrier. We also assume that blood flow has no influence on the probability of exchange occurring. In all cases it is possible to extend the generative model to account for these additional possibilities, although incorporation into the MRF framework would pose substantial computational challenges.

### 4.2 Healthy Volunteer Testing

This is the first demonstration of MRF methods to provide regional mapping in the human brain of blood brain barrier water exchange. All parameters showed good agreement with literature ranges.^18,8,21,19,22^ Expected differences between white and grey matter *T*_1,*t*_ and *v*_*b*_ are observed, while no differences are seen in *τ*_*b*_ or *T*_1,*b*_. Blood *T*_1_ would be expected to be approximately the same in both tissues, although differences in mean oxygenation may lead to small differences. It is unclear from the existing literature if we would expect to see differences *τ*_*b*_ regionally, although systematic differences in capillary geometry between white and grey matter could be hypothesised to lead to decreased *τ*_*b*_ in GM due to the increased surface area of vessels^46^. In addition, the maps show good agreement between hemispheres, suggesting physiologically relevant regional information is being extracted, rather than noise-based fluctuations. Little bias and generally good repeatability is observed in across regions. Quantification of grey matter parameters were less robust than the white matter parameters, which is likely due to the slice chosen including a relatively small volume of grey matter, leading to fewer voxels to average over. Grey matter is more likely to suffer from partial volume from white matter and CSF, which may explain the slightly low blood volume estimates.

The data presented here are regionally averaged, as the voxelwise data are noisy. However, it is possible that denoising strategies may mitigate this impact. In addition, when the error in the matching is evaluated (Supplementary Information Figure 3), it can be seen that noise in the images contains a strong coherent component that predominantly affects the anterior right of the brain surrounding the ventricles. This may be attributed to a previously identified issue with ordering of the spiral interleaves during reconstruction^47^ and CSF pulsation. The coherent nature of this artefact causes bias in the data, disrupting matching. Implementation of a modified reconstruction or denoising technique could be applied to alleviate this issue.

A limitation of implementing this work in clinical populations is the length of the acquisition, at 19 minutes, and in this work only a single slice is acquired due to limitations in the base MRF sequence that the protocol was constructed with. However, due to the short *TE* and long *TR*s used, multi slice acquisition could be implemented in this time.^48,49^ Acceleration strategies, such as in-place parallel reconstruction and simultaneous multi-slice (SMS) methods may also prove to be beneficial: SMS would allow for multiple slices to be acquired in the same acquisition time.

A key limitation of using MRF for the quantification of any parameter is the inherently discrete nature of the quantification. The dictionary to which the experimental data is matched is finite, and due to computational constraints often modest, which leads to discretisation errors in the final maps. This problem is a topic of much interest in the field, particularly in high dimensional problems such as this one. Application of a machine learning algorithm is often used to alleviate this issue^50,51,52^ and may be of benefit to our proposed approach.

## 5 CONCLUSIONS

Our simulations suggest that it is possible to optimise an SGRE-MRF acquisition for the quantification of blood volume and residence time of water in blood, simultaneously with *T*_1,*b*_, *T*_1,*t*_, and 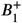. Our initial findings in healthy volunteers suggest that it is feasible to regionally quantify cerebral blood volume (*v*_*b*_) and mean intravascular residence time (*τ*_*b*_), simultaneously with *T*_1,*b*_, *T*_1,*t*_, and 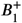 using an SGRE-MRF acquisition within a reasonable scan time. Our approach holds promise for patient-based measurements of subtle blood-brain barrier disruption.

## Supporting information

Supplementary Manuscript

## ACKNOWLEDGMENTS

This work is supported by the EPSRC-funded UCL Centre for Doctoral Training in Intelligent, Integrated Imaging in Healthcare (i4health) (EP/S021930/1), EPSRC grant EP/S031510/1, and the Department of Health’s NIHR-funded Biomedical Research Centre at University College London Hospitals. CWK receive funds from Horizon2020 (Research and Innovation Action Grants Human Brain Project 945539 (SGA3)), BRC (BRC704/CAP/CGW), MRC (MR/S026088/1), Ataxia UK, Rosetrees Trust (PGL22/100041 and PGL21/10079). The authors wish to thank Dr. David Higgins and Julia Markus for their help implementing this work in vivo.

## Financial disclosure

None reported.

## Conflict of interest

Geoff JM Parker declares the following potential conflicts of interests in companies with an interest in imaging biomarkers: Director and shareholder in Bioxydyn Limited; Director and shareholder in Quantitative Imaging Limited; Director and shareholder in Queen Square Analytics Limited.

Claudia AM Gandini Wheeler-Kingshott is a shareholder in Queen Square Analytics Limited.

## Notes

https://github.com/EmmaThomson/MRFSGRE_BBB

