## Supplementary Manuscript for "Quantification of water exchange across the blood-brain barrier using non-contrast MR fingerprinting"

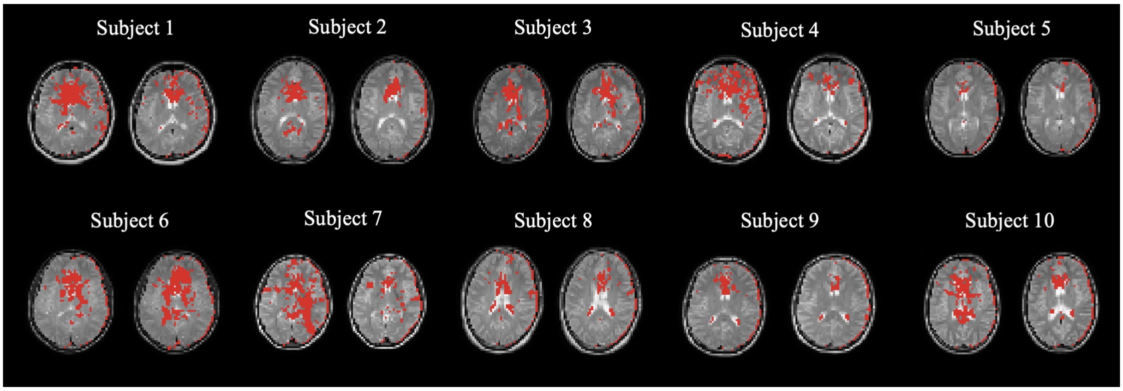


**FIGURE 1** Red voxels represent areas of low SNR that will result in a fit failure for $\tau_b$ and are therefore removed from calculations for all volunteers.

For P

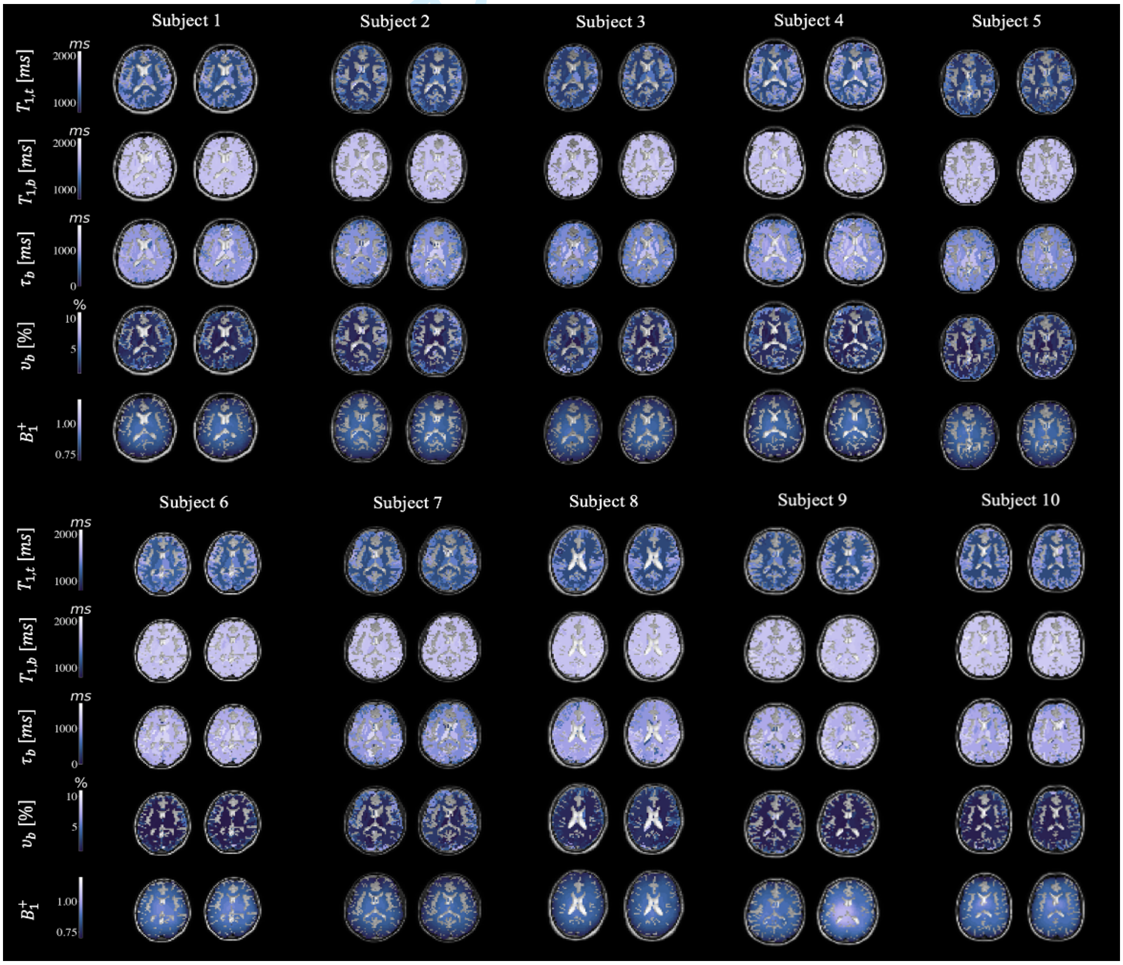


**FIGURE 2** Regional quantitative parameter maps of $T_{1,t}$, $T_{1,b}$, $v_b$ and $\tau_b$ for all volunteers with repeat scan, and a voxel wise map for $B_1^+$.

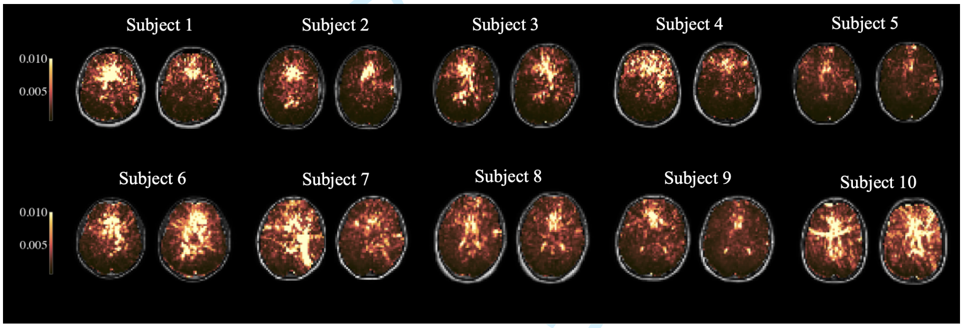


**FIGURE 3** Voxel wise maps of match quality of match for all volunteers represented by the residual sum of squares between the experimental signal and its closest dictionary match. Areas with a larger RSS are noisier.

For Peer Review
